# Optimization of interferential stimulation of the human brain with electrode arrays

**DOI:** 10.1101/783423

**Authors:** Yu Huang, Abhishek Datta, Lucas C. Parra

## Abstract

**Objective:** Interferential stimulation (IFS) has generated considerable interest recently because of its potential to achieve focal electric fields in deep brain areas with transcranial currents. Conventionally, IFS applies sinusoidal currents through two electrode pairs with close-by frequencies. Here we propose to use an array of electrodes instead of just two electrode pairs; and to use algorithmic optimization to identify the currents required at each electrode to target a desired location in the brain.

**Approach:** We formulate rigorous optimization criteria for IFS to achieve either maximal modulation-depth or maximally focal stimulation. We find the solution for optimal modulation-depth analytically and maximize for focal stimulation numerically.

**Main results:** Maximal modulation is achieved when IFS equals conventional high-definition multi-electrode transcranial electrical stimulation (HD-TES) with a modulated current source. This optimal solution can be found directly from a current-flow model, i.e. the “lead field” without the need for algorithmic optimization. Once currents are optimized numerically to achieve optimal focal stimulation, we find that IFS can indeed be more focal than conventional HD-TES, both at the cortical surface and deep in the brain. Generally, however, stimulation intensity of IFS is weak and the locus of highest intensity does not match the locus of highest modulation.

**Significance:** This proof-of-principle study shows the potential of IFS over HD-TES for focal non-invasive deep brain stimulation. Future work will be needed to improve on intensity of stimulation and convergence of the optimization procedure.

## 1. Introduction

Transcranial electric stimulation (TES) delivers weak electric current (≤ 2 mA) to the scalp in order to modulate neural activity of the brain (Nitsche and Paulus, 2000). Research has shown that TES can improve performance in some learning tasks and has also shown promise as a therapy for a number of neurological disorders such as depression, fibromyalgia and stroke (Nitsche et al., 2003; Fregni et al., 2006; Bikson et al., 2008; Schlaug et al., 2008). Conventional TES uses two sponge-electrodes to deliver current to the scalp. Modeling studies suggest that electric fields generated with such an approach are diffuse in the brain and typically cannot reach deep targets in the brain (Datta et al., 2009). A number of investigators have advocated the use of several small “high-definition” electrodes to achieve more focal or more intense stimulation (Datta et al., 2009; Dmochowski et al., 2011). This multi-electrode approach is referred to as high-definition transcranial electric stimulation (HD-TES, Edwards et al., 2013), and can be combined with any desired stimulation waveform.

Recently Grossman et al. (2017) proposed to stimulate the brain with “interferential stimulation” (IFS), which is well known in physical therapy (Goats, 1990). IFS applies sinusoidal waveforms of similar frequency through two electrode pairs. The investigators suggest that interference of these two waveforms can result in modulation of oscillating electric fields deep inside the brain. This promise of non-invasive deep brain stimulation has caused considerable excitement in the research community and in the news media (Dmochowski and Bikson, 2017; Shen, 2018). Our subsequent computational analysis of IFS suggests that it may not differ much from other multi-electrode methods in terms of intensity of stimulation (Huang and Parra, 2019). However, this conclusion is limited to the case of IFS using only two electrodes for each of the two frequencies. More recently researchers have turned to optimizing IFS in the hope that stimulation can be more focal in deep brain areas. A series of approaches have been proposed, but all are limited in some respect. Rampersad et al. (2019) searches through a large number of montages with two pairs of electrodes, but does not systematically tackle the general problem of arrays. Cao and Grover (2019) suggests the use of arrays, but fails to take interference into account by optimizing each frequency separately. Xiao et al. (2019) also proposes multiple pairs of electrodes but does not systematically optimize how to place them and what current to use through each.

Here we present a mathematical formulation of the optimization of IFS, including for the case of more than two electrode pairs. This allows us to systematically optimize the location of electrodes and the strength of injected currents through each electrode. This version of IFS using arrays differs from HD-TES only in that IFS uses two sinusoidal frequencies, whereas HD-TES uses the same waveform in all electrodes of the array. We show mathematically, and confirm numerically, that maximal modulation of interfering waveforms can be simply obtained by maximizing intensity of conventional HD-TES. We also provide a mathematical proof that the solution to maximum-intensity stimulation can be obtained directly from the forward model for TES, i.e. the “lead field” as first proposed by Fernandez-Corazza et al. (2019). Motivated by Guler et al. (2016) and Fernandez-Corazza et al. (2019), we convert the computationally intractable maximum-focality optimization for IFS into a maximum-intensity problem with constraint on the energy in the off-target area. This non-convex optimization is treated as a goal attainment problem (Gembicki and Haimes, 1975) and then solved with sequential quadratic programming (Brayton et al., 1979). IFS optimization in terms of intensity and focality is evaluated on the MNI-152 head model (Grabner et al., 2006) and compared to HD-TES solutions. Results show that one can achieve improved focality of modulation in deep brain areas as compared to HD-TES. This opens the possibility for non-invasive focal deep brain stimulation in the future.

## 2. Methods

### 2.1. Mathematical formulation

The mathematical framework for optimizing arrays of electrodes for TES was first introduced by Dmochowski et al. (2011). An important conclusion of that work was that there is a fundamental trade-off between achieving intense stimulation on a desired target, versus achieving focally constrained stimulation at that location. Thus, a number of competing optimization criteria were proposed in that work. Since then additional criteria have been proposed that strike a different balance or improve on computational efficiency. The criterion we used here is heavily motivated by Guler et al. (2016) because their formulation readily extends to IFS. Importantly, we now realize that the formulation allows one to readily adjust the trade-off between intensity and focality of stimulation. Let us explain.

The goal of targeting is to find an optimal source distribution **s** across a set of candidate electrode locations on the scalp. With *M* electrode locations this vector **s** has *M* dimensions, including one location reserved for the reference electrode. Since all currents entering the tissue must also exit, the sum of currents has to vanish,

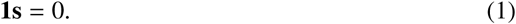

Here **1** is a *M*-dimensional row vector with all values set to 1. The inner product with this vector executes a sum. The electric field generated in the tissue by this current distribution **s** on the scalp is given by:

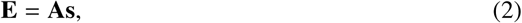

where **A** is the “forward model” for TES, also known as the “lead field” in EEG source modeling (Dmochowski et al., 2017). Specifically, it is a matrix that quantifies in each column the electric field generated in the tissue when passing a unit current from one electrode to the reference electrode. If there are *M* electrode locations on the scalp and *N* locations in the brain to be considered, then matrix **A** has *M* columns and 3*N* rows (one for each of the three directions of the field). The column corresponding to the reference electrode has all zero values as current enters and exits through the same electrode, i.e. no current flows through the tissue, and no field is generated. Note that this column is included in matrix **A** here to simplify all the equations throughout this paper. In the actual implementation matrix **A** does not have this all-zero column.

One goal of the optimization may be to maximize the intensity of stimulation at the target in a desired direction:

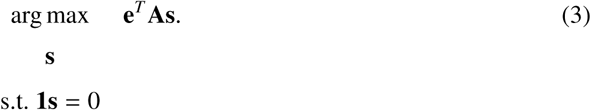

Here **e** is a column vector with 3*N* entries specifying the desired field distribution and direction at the target. For example, the values of **e** might be set to zero everywhere, except at the target location. If the desired target covers more than one location, then the non-zero values extend over all the corresponding locations. The specific values of **e** determine the desired orientation of the electric field.

At a minimum this optimization should be subject to the constraint that the total current injected shall not exceed a maximum value *I*_*max*_, otherwise this might cause discomfort or irritation on the scalp. The total applied current is the sum of absolute current through all electrodes:

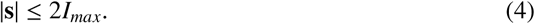

Here the notation | · | indicates the L1-norm, i.e., 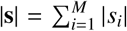. As the total current has to enter and exit the head it contributes to this sum twice, which explains the factor of 2 in the upper bound. In addition to this total current constraint, Guler et al. (2016) proposes to constrain the power of the electric field in brain areas outside the target region:

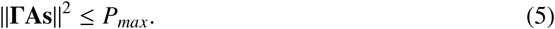

Here ‖ · ‖ means the L2-norm. The spatial selectivity for the off-target area is captured here by the diagonal matrix Γ. The diagonal elements of this matrix are zero for the target region, and non-zero elsewhere. These non-zero values can additionally implement a relative weighting for different locations (say, to compensate for uneven sampling in the FEM mesh, or to emphasize some regions more than others, although we do not do this here).

A few comments are in order here:

As pointed out by Guler et al. (2016), this quadratic criterion can be evaluated efficiently as, **s**^*T*^ **Qs** ≤ *P*_*max*_, which is fast to compute because, **Q** = **A**^*T*^ **Γ**^2^**A**, is a compact *M* × *M* matrix.

Constraints (4) and (5) specify a convex feasibility region and thus the optimization of the linear criterion can be solved with convex programming (see Section 2.5). In fact, when only the linear constraint (4) is active the solution can be found without the need for numerical optimization at all. We provide a prescription for how to find these solutions, along with a proof in Section 2.3.

The power constraint (5) can be used to titrate between maximally intense stimulation and maximally focal stimulation. Specifically, if *P*_*max*_ is set very high so as to not limit currents at all, then we are left only with the constraint (4) on current intensity. Criterion (3) subject to constraint (4) is the optimization proposed by Dmochowski et al. (2011) to achieve maximally intense stimulation. If, on the other hand, *P*_*max*_ is set to a very stringent value, then the constraint (5) dominates the maximization problem as shown by Guler et al. (2016). In that case we are left with criterion (3) subject to the power constraint (5). Fernandez-Corazza et al. (2019) show that this is equivalent to the least-squares criterion, which was introduced by Dmochowski et al.(2011) to achieve focal stimulation. The equivalence is only approximate, but becomes accurate if the target region is chosen to be small. In Section 3.1 we will show how intermediate values of *P*_*max*_ can move us between these two extremes of maximal intensity versus maximal focality.

Now we move on to the issue of optimizing IFS with electrode arrays.

### 2.2. Optimization of intensity in interferential stimulation

In inferential stimulation two fields oscillating sinusoidally at similar frequencies interfere to cause an amplitude modulation of the oscillating waveform. As an illustrating example, we show two sinusoidal electric fields **E**_**1**_ and **E**_**2**_ with frequencies *f*_1_ = 10 Hz and *f*_2_ = 11 Hz in Figure 1AB (red and blue curves). These two interfere to generate a modulated waveform referred to as the carrier (Figure 1CD, black curve). The envelope (green) of the modulated carrier oscillates at the difference frequency Δ*f* = 1 Hz. We can define two quantities: (1) intensity of the carrier signal – the peak-to-zero amplitude of the black curve in Figure 1CD; (2) modulation depth – the peak-to-peak amplitude of the green curve in Figure 1CD, which is 2 min(‖**E**_**1**_‖, ‖**E**_**2**_‖) (Huang and Parra, 2019). In Figure 1 we see how the amplitudes of the two interfering signals in IFS affect the modulation depth: in panel A the two signals have different amplitudes, leading to a carrier waveform of 2 V/m in panel C and a modulation depth of 1 V/m (i.e., 50% modulation); when the two interfering signals have the same amplitudes (panel B), we see the carrier intensity and modulation depth are both 2 V/m (i.e., 100% modulation, panel D). In fact, HD-TES can apply a single waveform of any shape, in particular, it can just apply a 100% modulated sinusoid (black curve in panel D) to ensure the modulation depth is the same as the carrier intensity. On the other hand, modulation depth is weaker than signal intensity in IFS if the two interfering currents have different amplitudes. We use the terms “modulation depth” and “intensity” interchangeably in this paper but we only optimize the modulation depth for both HD-TES and IFS (see Figures 3, 4 and 5 for the differences).

**Figure 1:**
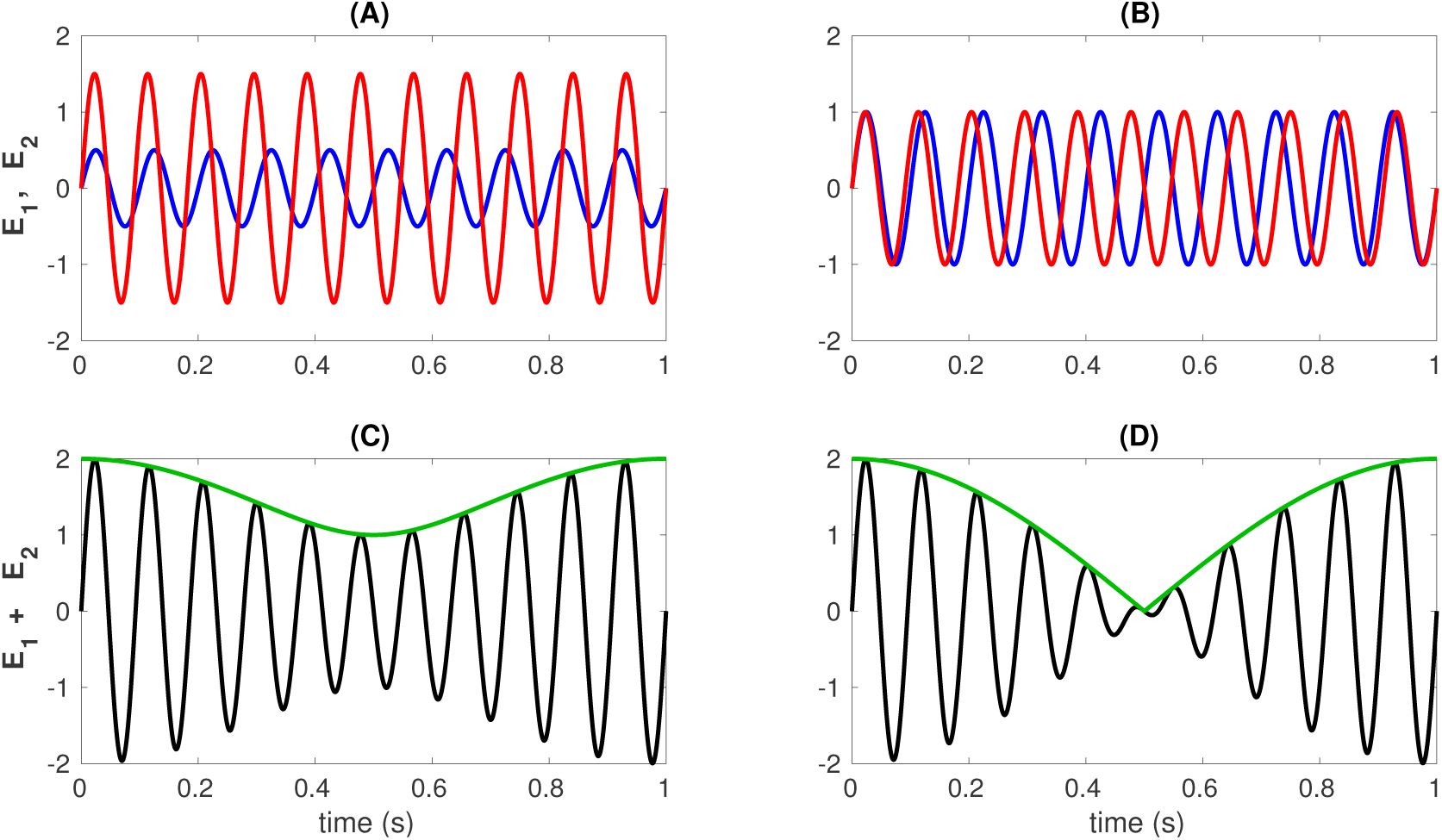
Modulation vs. Intensity: (AB) sinusoidal electric fields **E**_**1**_ and **E**_**2**_ with frequencies *f*_1_ = 10 Hz and *f*_2_ = 11 Hz (red and blue) in IFS. (CD) **E**_**1**_ and **E**_**2**_ interfere to generate a modulated waveform known as the carrier (black). The envelope (green) of the modulated carrier oscillates at the difference frequency Δ*f* = 1 Hz. For IFS modulation depth can only be 100% when both fields at a given location have the same magnitude (BD). In most locations fields will not be the same for both frequencies and modulation depth will be less than the intensity (AC). The only way for both fields to be identical everywhere for IFS is for current amplitude to be identical for both frequencies in all electrodes. HD-TES can simply apply a single waveform of 100% modulated sinusoid (black curve in panel D).

**Figure 2:**
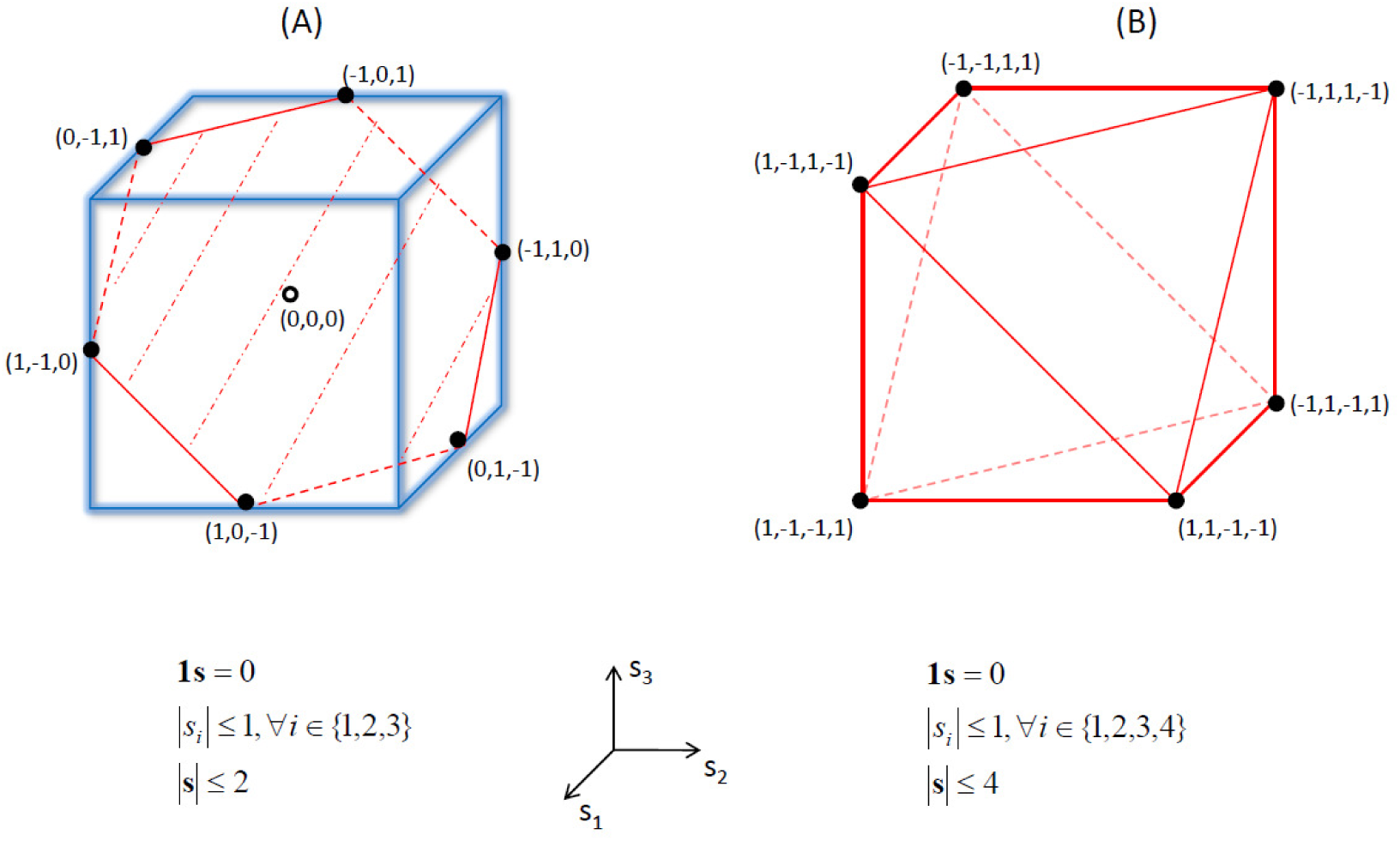
Linear constraints define a polyhedral feasibility region. With a linear criterion, possible solutions are found at the corner points (black dots). (A) Two-dimensional feasibility region for *M* = 3 electrodes. The blue cubic bounding box represents the current constraint at each electrode (10), with *I*_*max*_ = 1. This constraint is intersected by the plane defined by the zero-sum-current constraint (1). This results in the hexagonal feasibility region (shaded red).(B) Three- dimensional feasibility region for *M* = 4 electrodes. Here the constraints on the total current and the constraint at each electrodes do not coincide.

**Figure 3:**
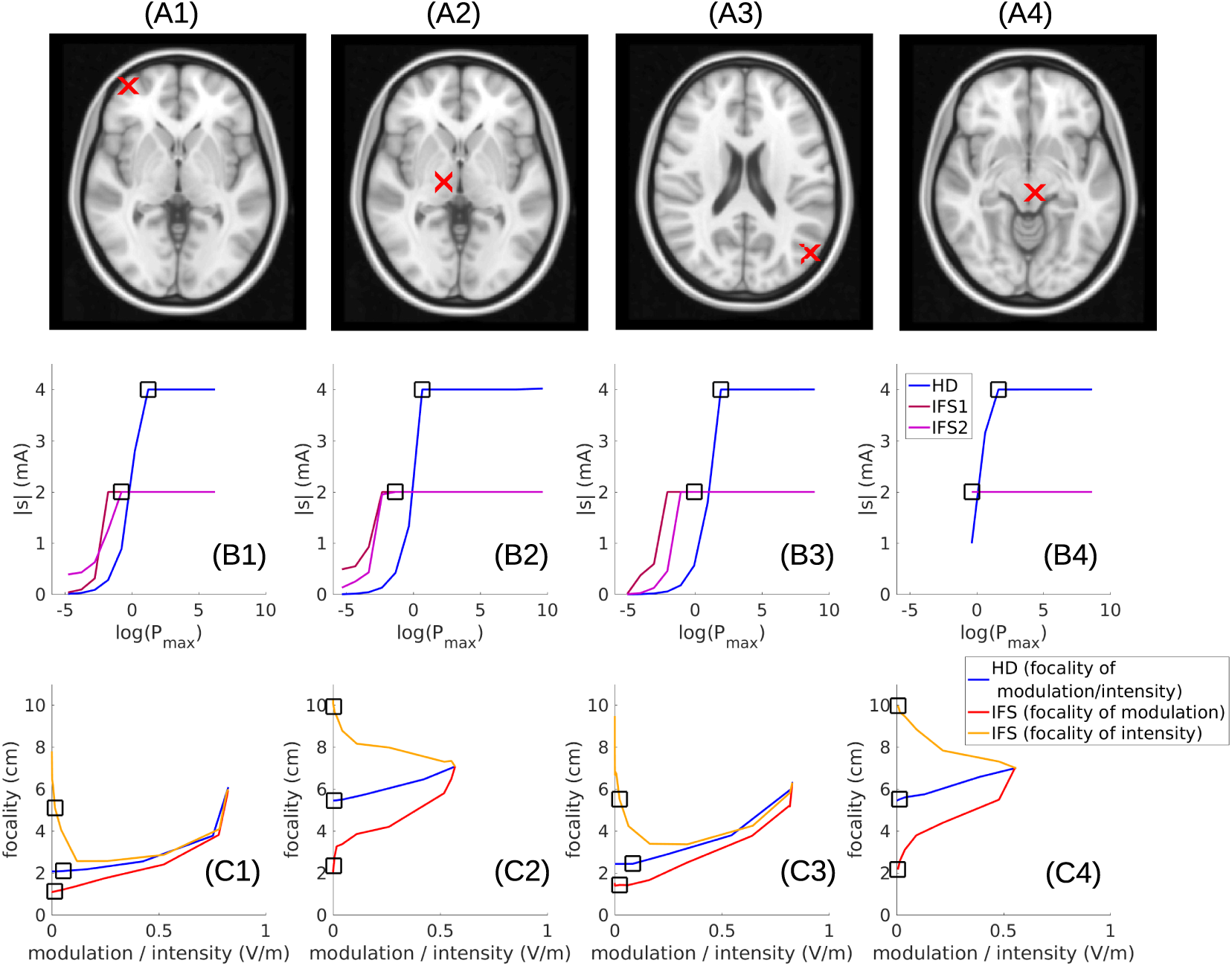
Optimal field at the target for varying power constraints at off-target areas. Target locations are indicated by the red x in the MRI, with 2 on the cortical surface (panels A1, A3), and 2 in the deep brain (panels A2, A4). Desired field orientation is in the radial direction. (B1–B4) Total injected current intensity (|**s**|) as the function of the power constraint *P*_*max*_ on a logarithmic scale. IFS1 and IFS2 are the total current injected at the two frequencies of IFS, each going to a maximum of 2mA. For HD the maximum is 4mA. (C1–C4) Intensity versus focality for different values of *P*_*max*_, with squares indicating injected current reaches the safety limit. Note for IFS, both modulation depth and intensity are shown.

**Figure 4:**
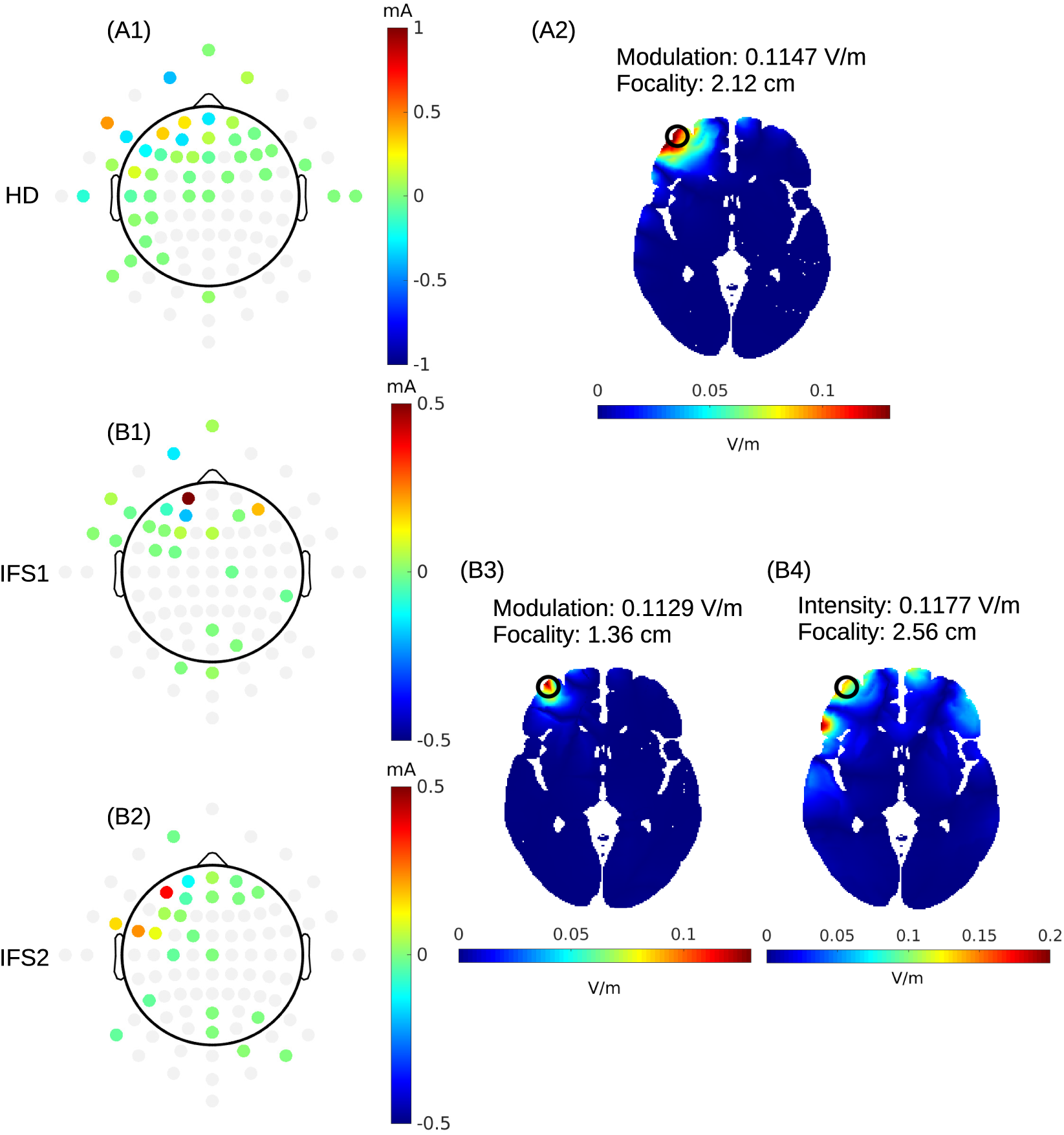
Example of maximal modulation depth under strong off-target power constraints. These solutions are selected to have a comparable maximum intensity for HD-TES (panels A) and modulation depth for IFS (panels B). A1: Current distribution across HD-TES electrodes. B1 & B2: current distribution for IFS at the two frequencies. Gray dots indicate candidate electrodes with no current injected. Radial component of the modulation depth is shown for HD-TES (panel A2) and IFS (panel B3). The intensity of electric field is also shown for IFS (panel B4). The target is located at the cortical surface (black circles) and desired field points in radial direction. Modulation depth / intensity and focality are noted for the target location.

**Figure 5:**
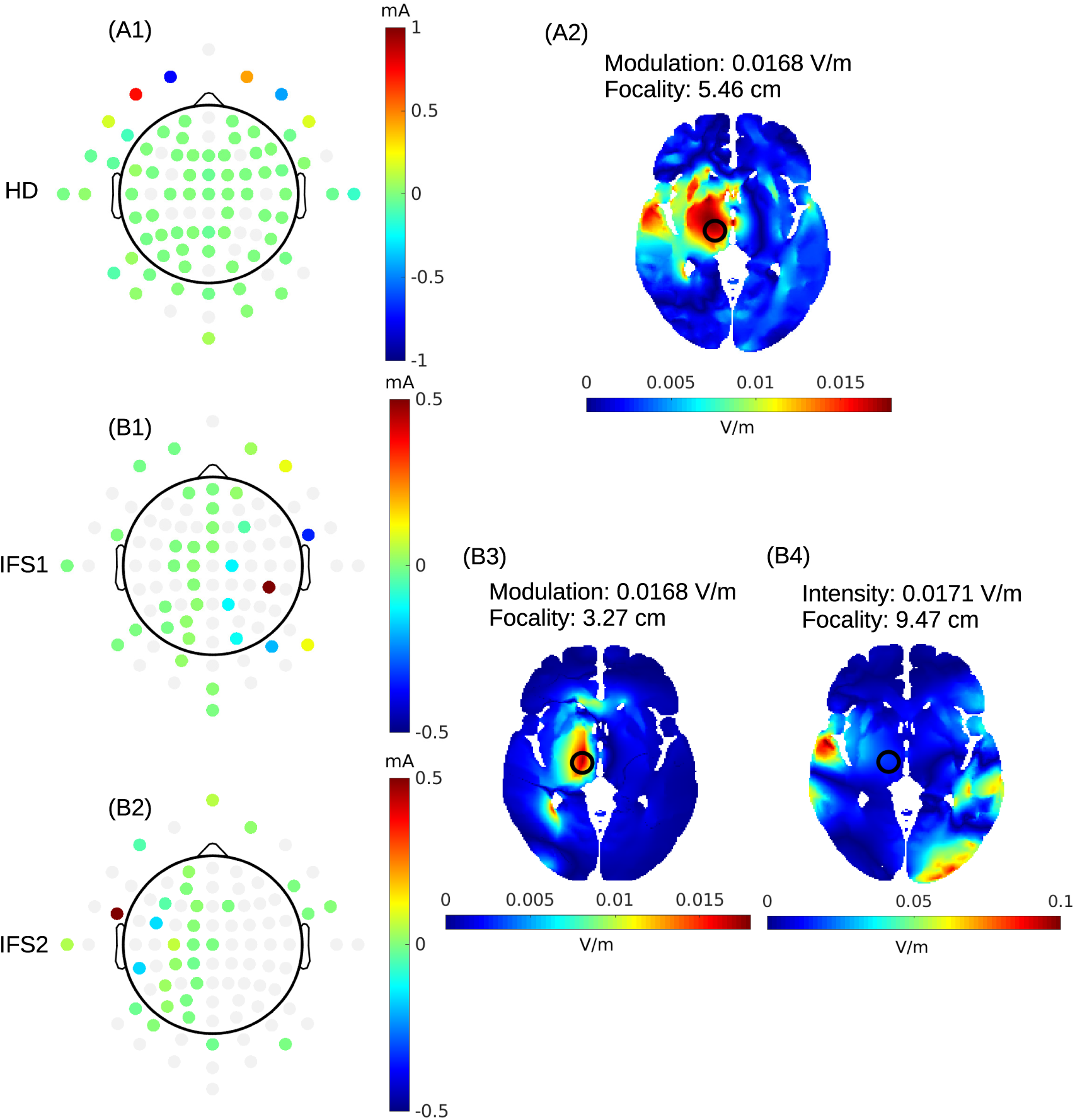
Same as Figure 4 but for a deep target (black circles).

For IFS, let’s assume that **E**_**1**_ and **E**_**2**_ are generated by the current distributions **s**_1_ and **s**_2_, each oscillating at their own frequency. To maximize the modulation depth 2 min(‖**E**_**1**_‖, ‖**E**_**2**_‖) at a location along an orientation both specified by vector **e**, we propose the following criterion:

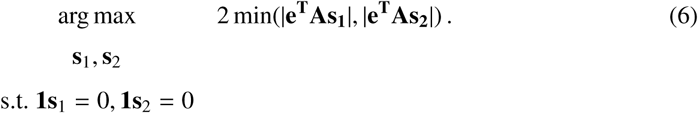

The vectors **s**_1_ and **s**_2_ are both of length *M*. The zero sum constraints are needed to maintain physical feasibility. The vectors quantify currents in the same set of electrodes. Thus, each single electrode has the freedom to pass current at two different frequencies and intensities added together. This generalizes the conventional IFS approach (Grossman et al., 2017) in that each frequency is now applied to potentially more than one electrode pair. It is also more flexible than recent efforts to target IFS with multiple pairs as they are limited to applying only one oscillating frequency at each pair (Rampersad et al., 2019; Cao and Grover, 2019). In contrast, here each electrode can apply the sum of two oscillating currents, and thus we can variably distributed the two frequencies over all electrodes in the array.

For the sake of safety and comfort we again limit the total applied current to a maximum value:

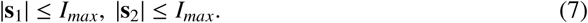

Here each current distribution is limited by *I*_*max*_, such that the total injected current is bounded by the same value as in HD-TES (Eq. 4).

Note that the optimization criterion (6) is bounded from above:

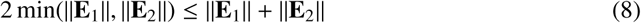

and the equality holds, if and only if ‖**E**_1_‖ = ‖**E**_2_‖. This implies that no matter what choice one makes for **s**_1_, the criterion (6) will always be maximal for **s**_2_ = **s**_1_ as then ‖**E**_1_‖ = ‖**E**_2_‖ (Figure 1BD). In other words, we can equivalently optimize for a single **s** = **s**_1_ = **s**_2_ (subject to the same set of constraint):

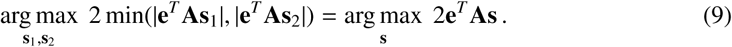

This criterion (9) is the same maximum-intensity criterion as (3), which we used for HD-TES. What does it mean that currents through electrodes are the same for both frequencies? It simply means that each electrode is passing the same amplitude-modulated waveform (with 100% modulation depth) with magnitude and sign as defined by **s**. Therefore, to optimize the modulation depth of IFS one simply needs to “fuse” the two current sources. In short, the largest modulation depth of IFS will be achieved with the max-intensity solution of HD-TES. In Section 3.2 we provide numerical confirmation for this theoretical result, namely, that the maximum intensity criterion gives the same result for IFS and HD-TES.

### 2.3. Closed form solution for max-intensity optimization

For the maximum-intensity problem of HD-TES (Eq. (3) subject to (4)), it is sufficient to write the maximum total current constraint (4) as a constraint on each electrode, when taking into account that in both cases the sum of all currents has to be zero as in (1):

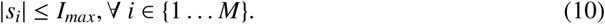

Linear inequalities such as (10) & (1) specify a convex polyhedron, see e.g., Figure 2A, a two-dimensional polyhedron for *M* = 3 electrodes and *I*_*max*_ = 1. With a linear optimality criterion this constitutes a linear programming problem (Grant et al., 2006). The fundamental theorem of linear programming states that the solutions are at the corner of the polyhedral feasibility region (black dots in Figure 2). For the example in Figure 2A the corners correspond to current *I*_*max*_ passing through a single pair of electrodes *i, j* that gives the largest value for **e**^*T*^ **As**. If we define vector

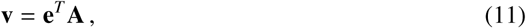

then *i* is simply the position with the largest value in **v**, and *j* is the position with the smallest value. Therefore, the optimal solution can be found directly from the forward model **A**. This has been first noted by Fernandez-Corazza et al. (2016) and is discussed in Fernandez-Corazza et al. (2019) and Saturnino et al. (2019). We could have also relaxed the total current constraint by allowing twice as much current compared to the limit on each electrode, |*s*_*i*_| ≤ *I*_*max*_. This changes (4) to |**s**| ≤ 4*I*_*max*_. Now the total current has to be split among at least two electrode pairs (four electrodes total), see e.g., Figure 2B, a three-dimensional feasibility region for *M* = 4 electrodes with *I*_*max*_ = 1. The maximum intensity on target will be achieved if we inject +*I*_*max*_ though the two electrode locations with the largest values of **v** and draw -*I*_*max*_ at the two locations with the smallest values of **v**. Of course this argument applies not just for two pairs of electrodes, but any number of electrodes needed to achieve a larger allowable total current. The purpose of a lower limit on individual electrodes is to distribute a larger current over multiple locations while limiting sensation at each electrode. We have done this in previous work and found the solutions with numerical optimization (Dmochowski et al., 2011; Huang and Parra, 2019). Now we realize that finding the optimal solution does not require any numerical optimization as proposed in Dmochowski et al. (2011).

### 2.4. Optimization of focality in interferential stimulation

To optimize interferential stimulation in terms of focality we add the constraint on the amplitude modulation at off-target locations. In analogy with (5) we constrain the square of modulation depth, summed over all locations in the off-target region:

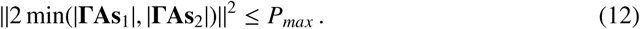

Note here the absolute value is taken element-wise for **ΓAs**_1_ and **ΓAs**_2_. This is a non-convex constraint which complicates the numerical optimization. We solved optimization of the IFS criterion (6) s.t. (7) and (12) by sequential quadratic programming (SQP) (Brayton et al., 1979), which is implemented in the “`fminimax`” function in Matlab (R2016, MathWorks, Natick, MA).

### 2.5. Implementation details

To solve the convex optimization problems, such as for HD-TES (3 s.t. 4, 5) we use the CVX toolbox (Grant et al., 2006). This Matlab toolbox provides a high-level language interface that allows one to specify L1 norm constraints such as (4). Unfortunately it does not implement the minimum operation in (6). For this we introduce an auxiliary variable *z* and solve equivalently:

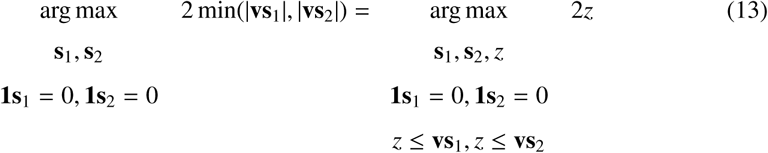

To solve the non-convex optimization for IFS (6 s.t. 7, 12) we use the “`fminimax`” function in Matlab. This function allows one to directly implement the minimum operation in (6). Unfortunately it does not directly implement the L1-norm constraint (7). Following the method proposed in Tibshirani (1996) we can formulate this constraint as set of simple linear constraints using auxiliary variables **s**^+^ and **s**^-^, which substitute for variable **s**:

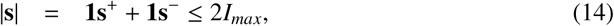

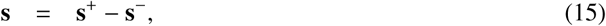

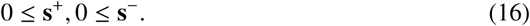

When adjusting the power constraint *P*_*max*_ at off-target area, we first calculate a default value of *P*_*max*_ as **e**^*T*^ **A**(**A**^*T*^ **Γ**^2^**A**)^-1^**A**^*T*^ **e**, which is the value that makes the criterion equivalent to least-squares criterion for maximum-focality (Fernandez-Corazza et al., 2019). We then vary this value across different orders of magnitude: for superficial targets (Figure 3, A1&A3), we vary the power constraint from *P*_*max*_ ×10^−3^ to *P*_*max*_ ×10^8^; for deep targets (Figure 3, A2&A4), we relax the power constraint furthermore to *P*_*max*_ × 10^12^ as the optimization is numerically unstable for deep targets when the power constraint is very stringent. When solving the IFS (6 s.t. 7, 12) for the first value of power constraint (i.e., *P*_*max*_ ×10^−3^), we initialize the “`fminimax`” function using the solution from HD-TES under the same power constraint (see Supplementary Information for why we need to do this). Then as the power constraint increases (relaxes), each solving for the IFS is initialized by the previous solution. For each value of *P*_*max*_, the modulation depth / intensity and focality of the optimized electric field at the target location are computed. The focality is defined as the cubic-root of the brain volume with electric field modulation depth / intensity of at least 50% of the field intensity at the target (Huang and Parra, 2019).

In summary, to optimize the modulation depth of IFS, we implemented in the CVX toolbox these equations:

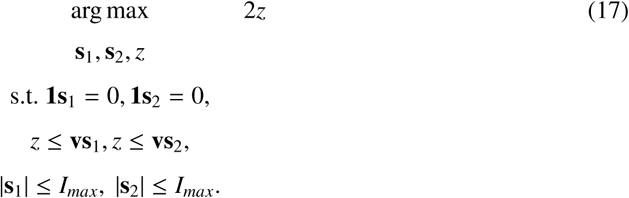

To optimize the focality of IFS, we implemented the following equations using the “`fminimax`” function in Matlab, with varying values of *P*_*max*_:

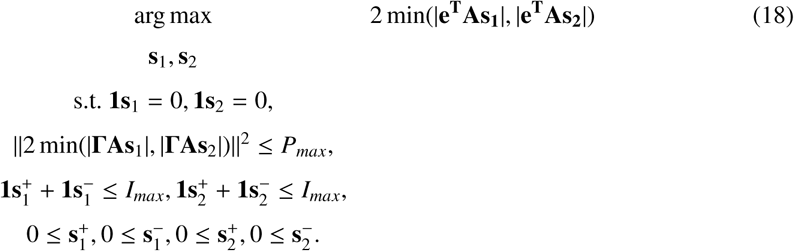

Note here 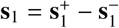 and 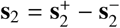.

### 2.6. Construction of the head model

The forward model **A** in this work was built on the ICBM152 (v6) template from the Montreal Neurological Institute (MNI, Montreal, Canada) (Mazziotta et al., 2001; Grabner et al., 2006)), following previously published routine (Huang et al., 2013). Briefly, the ICBM152 (v6) template MRI (magnetic resonance image) was segmented by the New Segment toolbox (Ashburner and Friston, 2005) in Statistical Parametric Mapping 8 (SPM8, Wellcome Trust Centre for Neuroimaging, London, UK) implemented in Matlab. Segmentation errors such as discontinuities in CSF and noisy voxels were corrected first by a customized Matlab script (Huang et al., 2013) and then by hand in an interactive segmentation software ScanIP (v4.2, Simpleware Ltd, Exeter, UK). Since tDCS modeling work has demonstrated the need to include the entire head down to the neck for realistic current flow, in particular in deep-brain areas and the brainstem (Huang et al., 2013), the field of view (FOV) of the ICBM152 (v6) MRI was extended down to the neck by registering and reslicing the standard head published in (Huang et al., 2013) to the voxel space of ICBM152 (see Huang et al. (2016) for details). HD electrodes following the convention of the standard 10–10 international system (Klem et al., 1999) were placed on the scalp surface by custom Matlab script (Huang et al., 2013). Two rows of electrodes below the ears and four additional electrodes around the neck were also placed to allow for targeting of deeper cortical areas and for the use of distant reference electrodes in tDCS. A total of 93 electrodes were placed. A finite element model (FEM, (Logan, 2007)) was generated from the segmentation data by the ScanFE module in ScanIP. Laplace’s equation was then solved (Griffiths, 1999) in Abaqus 6.11 (SIMULIA, Providence, RI) for the electric field distribution in the head. With one fixed reference electrode Iz as cathode, the electric field was solved for all other 92 electrodes with unit current density injected for each of them, giving 92 solutions for electric field distribution representing the forward model of the ICBM152 head, i.e., the matrix **A**. Note this matrix **A** is also the transpose of the EEG lead field **L** (Dmochowski et al., 2017). **This data is available at https://www.parralab.org/optIFS/.**

## 3. Results

### 3.1. Off-target power controls focality versus intensity; IFS achieves more focal modulation depth

The method proposed here maximizes the modulation depth at the target while constraining the power in off-target areas (see Section 2.1, 2.2, and 2.4). Note that for IFS modulation depth is generally smaller than intensity (Figure 1). For HD-TES the two are identical so that maximizing intensity is equivalent to maximizing modulation depth. We first performed numerical experiments to determine how the power constraint affects the optimization results for both HD-TES and IFS. Specifically, we numerically solved for the HD-TES criterion (3 s.t. 4, 5) and the IFS criterion (6 s.t. 7, 12). The desired field direction **e** was selected to point in the radial-in direction, and we varied the maximum allowable off-target power *P*_*max*_. We computed the modulation depth, intensity and focality of the optimized electric field at the target location for different values of *P*_*max*_ (Figure 3, C1–C4; see Section 2.5 for details). This was done for four different target locations in the brain: 2 on the cortical surface and 2 in the deep brain region (Figure 3, A1–A4). Following the common safety standard, total current was selected to not exceed 2 mA (or an L1-norm of 4 mA;). For IFS, this is 1 mA (or an L1-norm of 2 mA) for each of the two frequencies. For small values of *P*_*max*_ the total currents do not reach this allowable maximum, indicating that the power constraint dominates (Figure 3, B1–B4, left of square markers). Note for deep target A4, we do not show the IFS results under small *P*_*max*_ as we found that the optimization is numerically unstable before the total injected current reaches the maximum. When the power constraint is relaxed beyond this threshold values, then the full current is used (right of square markers). As we had predicted based on theoretical grounds (in Section 2.1), *P*_*max*_ regulates the trade-off between intensity and focality (Figure 3, C1–C4). Generally, as the power constraint in the off-target region is relaxed, modulation depth increases and the area of stimulation increases in size. This trend is evident for both IFS and HD-TES. Again, note that for IFS modulation depth is different from the intensity (Figure 1), while these two are the same for HD-TES. Evidently, for the same modulation depth at the target, the optimized IFS provides more focal modulation compared to optimized HD-TES (red curve is below blue curve in Figure 3, C1–C4). The intensity of IFS, however, is less focal than either IFS modulation depth or HD intensity (orange curve is mostly higher than both red and blue in Figure 3, C1–C4). The advantage of optimized IFS compared to optimized HD-TES in terms of focal modulation is more evident in the deep targets than the superficial targets (bigger gap between red and blue curves in Figure 3 C2 & C4 than in C1 & C3. We also note that the modulation / intensity is generally weaker in the deep locations (up to 0.6 V/m, C2&C4) compared to the cortical locations (up to 0.8 V/m, C1&C3). Finally, when the off-target power constraint *P*_*max*_ is relaxed enough, IFS and HD-TES converge to the same results of maximal-intense stimulation (see Section 2.1 and 2.2, also Figure 6 and Supplementary Figures 2–5).

**Figure 6:**
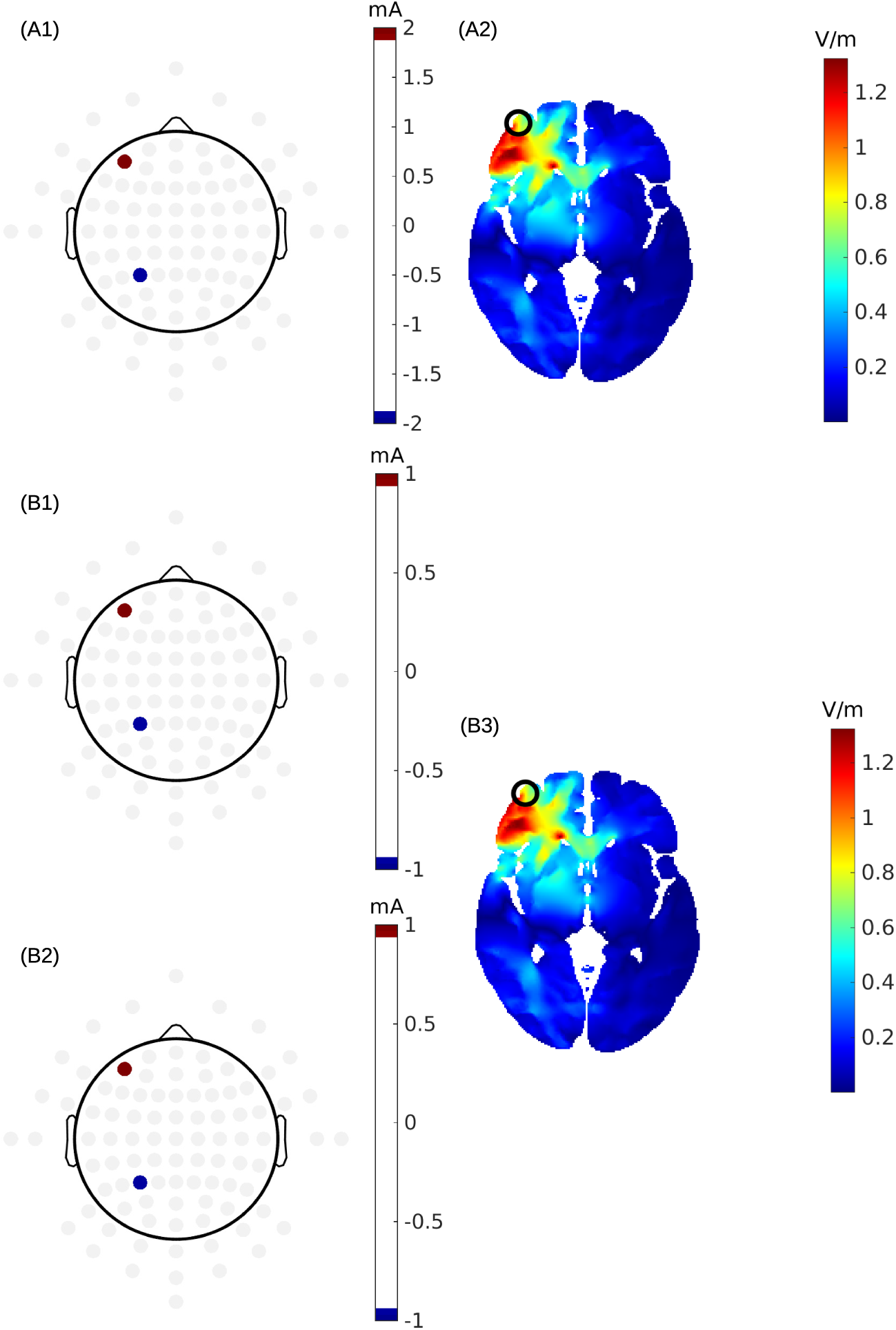
Numerical solutions of maximizing the electric field without an off-target power constraint for conventional HD-TES (panel A) and interferential stimulation (panel B). Gray dots in the electrode layouts indicate candidate electrodes with no current injected. The target location is indicated by a black circle in panels A2 and B3, where the radial component of the modulation depth is shown.

### 3.2. Numerical solutions on a few examples

For the first two target locations (Figure 3, A1 and A2) we fix *P*_*max*_ to the threshold value where current is fully utilized without losing focality (square points in Figure 3). The resulting optimized modulation depth and field intensity are shown in Figure 4 and Figure 5. The two montages in panels B1 and B2 correspond to the currents **s**_1_ and **s**_2_ with the two different frequencies, injected through the same array. For IFS, modulation depth is different from the intensity, as shown in panels B3 and B4, respectively. The modulation depth of IFS achieves a focal spot that is about 60% the size of that of HD-TES (1.36 cm vs. 2.12 cm at the cortical target, Figure 4; 3.27 cm vs. 5.46 cm at the deep target, Figure 5; all comparison under similar levels of modulation depth). However, the intensity of the electric field for IFS missed the target locations. Note the modulation depth in the deep brain (0.02 V/m, Figure 5) is much weaker than that on the cortical surface (0.11 V/m, Figure 4), and the modulation increases with the power constraint *P*_*max*_ relaxed, but at the price of losing focality (see Supplementary Information for more examples of these figures under different levels of *P*_*max*_).

When the power constraint *P*_*max*_ is removed (5 for HD-TES, 12 for IFS), we have the maximum-intensity solution. Examples of that are shown in Figure 6 for HD-TES (panel A) and IFS (panel B). As predicted mathematically (Section 2.2), one needs to inject the same current for the two frequencies at the same pair of electrodes. The location of these two electrodes can be simply determined as the largest and smallest values of vector **v**, which quantifies the voltages that would be generated by a current source at the target location in the brain (see Eq. 11). The resulting modulation depth in radial direction (panel B3) is exactly the same as the electric field along the same direction induced by the HD-TES (panel A2). Therefore, in terms of intensity on target, IFS can not be any stronger than conventional HD-TES, even when optimizing electrode placement, but IFS does gain focality in modulation depth when optimized.

## 4. Discussion and Conclusions

In this work we addressed the optimization of IFS by considering an array of electrodes that can apply sinusoidal currents at two different frequencies. Each electrode in the array can apply different current intensities for each frequency. This is significantly more flexible than recent optimization efforts for IFS that have been limited to pairs of electrodes of a single frequency each (Rampersad et al., 2019; Cao and Grover, 2019; Xiao et al., 2019). The approach can be readily implemented with existing current-controlled multi-channel TES hardware. The current sources with frequencies *f*_1_ and *f*_2_ are simply connected in parallel to the same electrodes.

We found that maximizing the modulation depth of IFS results in a solution that “fuses” the two frequencies, in the sense that they are to be applied with the same intensity in a given electrode. The IFS solution is then equivalent to HD-TES with a modulated waveform (with 100% modulation). We further prove that this maximum-intensity montage – which is optimal for both HD-TES and IFS – can be found directly from the forward model.

We previously established for HD-TES that there is a fundamental trade-off between focality and intensity of stimulation (Dmochowski et al., 2011). We find this trade-off here reproduced for the modulation depth of IFS. We leverage the optimization criterion of Guler et al. (2016) which allows one to titrate between the two extremes by constraining the power of stimulation in off-target areas: a tighter constraint makes the result more focal, a looser constraint results in stronger stimulation on target. When applying the same optimization criterion to the modulation depth of IFS we obtain a genuinely different optimization problem. The upside is that we find more focal stimulation for IFS as compared to HD-TES for otherwise similar modulation depth. This is true for cortical locations as well as deep brain areas. This appears to conflict with our previous conclusion on this topic (Huang and Parra, 2019), where we show that the two techniques were largely the same in terms of focality. One caveat of that earlier work was that the electrode arrays had not been systematically optimized, and thus we left the possibility open that IFS could be more focal than HD-TES. Here we conclude that this is in fact the case once currents through each electrode have been systematically optimized.

When designing stimulation montages, what should one focus on, intensity of the high frequency carrier, or its modulation depth? For HD-TES they can be readily made the same (Figure 1). For IFS they are not, and in fact as we show in Figures 4 and 5 they can be quite different. The premise of IFS is that high frequencies have no effect on nervous tissues, and instead, neurons and/or nerve fibers respond more readily to the modulation or transients of the high-frequency stimulation (Grossman et al., 2017). Thus, it may make sense to focus on modulation depth instead of intensity. However, empirical validation of this assumption is mixed (unpublished data). Therefore, while optimization of IFS has focused on the modulation depth here, we also report the associated intensities of the high frequency carrier.

The advantage in focality of IFS comes from the *min*() operation in calculating the modulation depth (Eq. 12). This operation is element-wise and it iterates over all the locations in the off-target area, which makes it computationally very expensive (optimization with the “fminmax” function takes 1–2 hours on a standard PC). Additionally, the optimization criterion becomes non-convex, which causes sub-optimal solutions if the search is not properly initialized (see Supplementary Figure 1). We have tested alternative criteria but find in all instances that the optimal solutions again “fuse” the two frequencies (see Appendix). In other words, without the location-wise *min*() operation we have not been able to achieve any different solution than the conventional HD-TES. The *min*() operation appears to break the symmetry of the solution leading to higher focality compared to HD-TES.

In conclusion, so far, the naive approaches advocated for IFS do not meaningfully outperform HD-TES. Yet, here we provide a proof-of-principle that more focal stimulation is possible with IFS. The challenge remains to find an efficient convex optimization criterion, and to bring intensity of stimulation and modulation depth into better agreement. Only then can one expect to exploit the full potential of IFS for the purpose of focal non-invasive deep brain stimulation.

## ACKNOWLEDGMENT

This work was supported by the NIH Grants R01MH111896, R44NS092144, R01NS095123, and by Soterix Medical Inc.

## Appendix

As an alternative to Eq. (12), one could limit an upper bound of the power, by exploiting the relationships: ‖ min(|**E**_1_|, |**E**_2_|)‖^2^ ≤ ‖**E**_1_‖^2^ + ‖**E**_2_‖^2^. With this we could limit the total power summed over off-target locations:

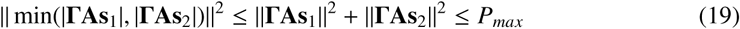

Alternatively, based on the upper bound (8), we could limit the amplitude at off-target locations:

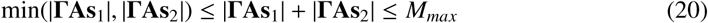

Both constraints (19) and (20) are convex so that the optimization of the linear criterion (3) subject to these constraints can be solved efficiently with established convex optimization tools (Section 2.5). We found numerically for both these constraints that the optimum is again given by the symmetric solution, **s** = **s**_1_ = **s**_2_, as in the max-intensity solution (Eq. 9).

## Supplementary Information

Here in this Supplementary Information, we show how initialization can affect the non-convex optimization of the focality of IFS. We also show the electric field distributions under different levels of off-target energy constraints.

### IFS optimization requires proper initialization

Finding the most focal IFS montage is a non-convex optimization problem (Eqs. 6 s.t. 7, 12). Thus the solution is subject to local minima, and the search depends on the initial conditions. We have compared the results of the search initialized with random current intensities, to solutions found by initialing with the optimal HD-TES results under the same off-target energy constraint. Specifically, for the first two targets (Figure 3, A1 and A2), we initialized the search using random currents following a standard Gaussian distribution with standard deviation of 1 mA. As the search is numerically expensive (“fminmax” function takes 1–2 hours on a standard PC), we only ran this 20 times for each target. For the superficial target (Figure S1 A1), we found that the objective function (Eq. 6) achieves similar levels following random initialization as compared to initialization with the HD-TES solution (blue bars vs. red line in Figure S1 B1). However, for the deep target (Figure S1 A2), a random initialization results in inferior performance (Figure S1 B2). Therefore, one should at least use the solution from the HD-TES optimization to initialize IFS optimization.

### Visualization of the electric field under different levels of off-target energy constraints

To give an intuition for the advantage of optimized IFS in terms of focal brain stimulation, we provide more examples of electric field distributions here under different levels of off-target energy constraints *P*_*max*_ (Figures S2 – S5). The advantage of focality in optimized IFS over optimized HD-TES is more obvious for deep targets (Figures S3 & S5) than for cortical targets (Figures S2 & S4). Also, we see that as *P*_*max*_ is relaxed, IFS and HD-TES tend to converge to the same results.

**Figure S1:**
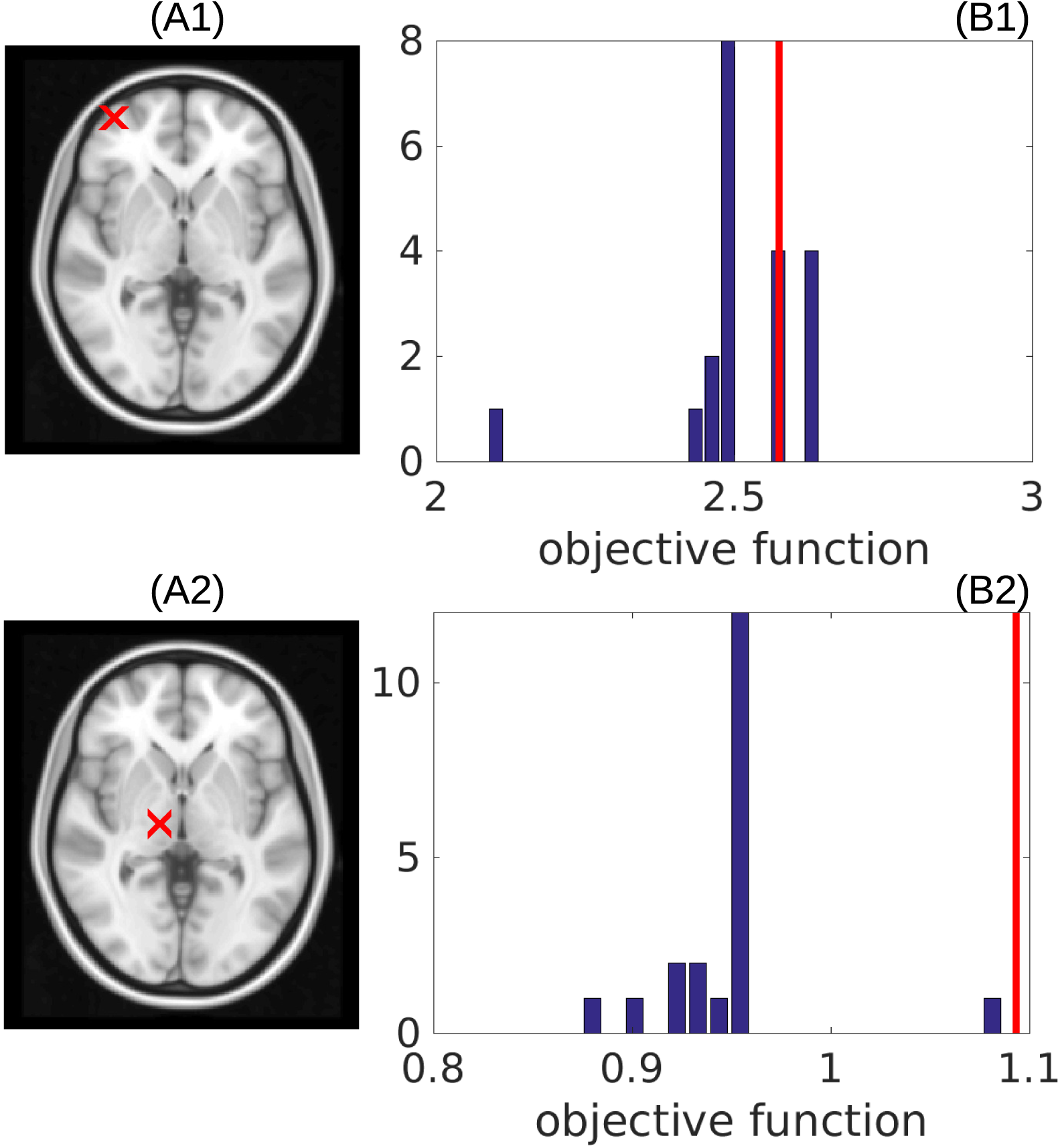
Effects of initialization for optimization of focality of IFS. Results are for a superficial target (A1) and deep target (A2), identical to Figure 4 and 5 respectively. Histograms of maximal values achieved for the objective function (Eq. 6) with random initialization are shown as blue bars in (B1) and (B2), with red vertical lines indicating the optimums obtained from initializing with the optimal HD-TES solution.

**Figure S2:**
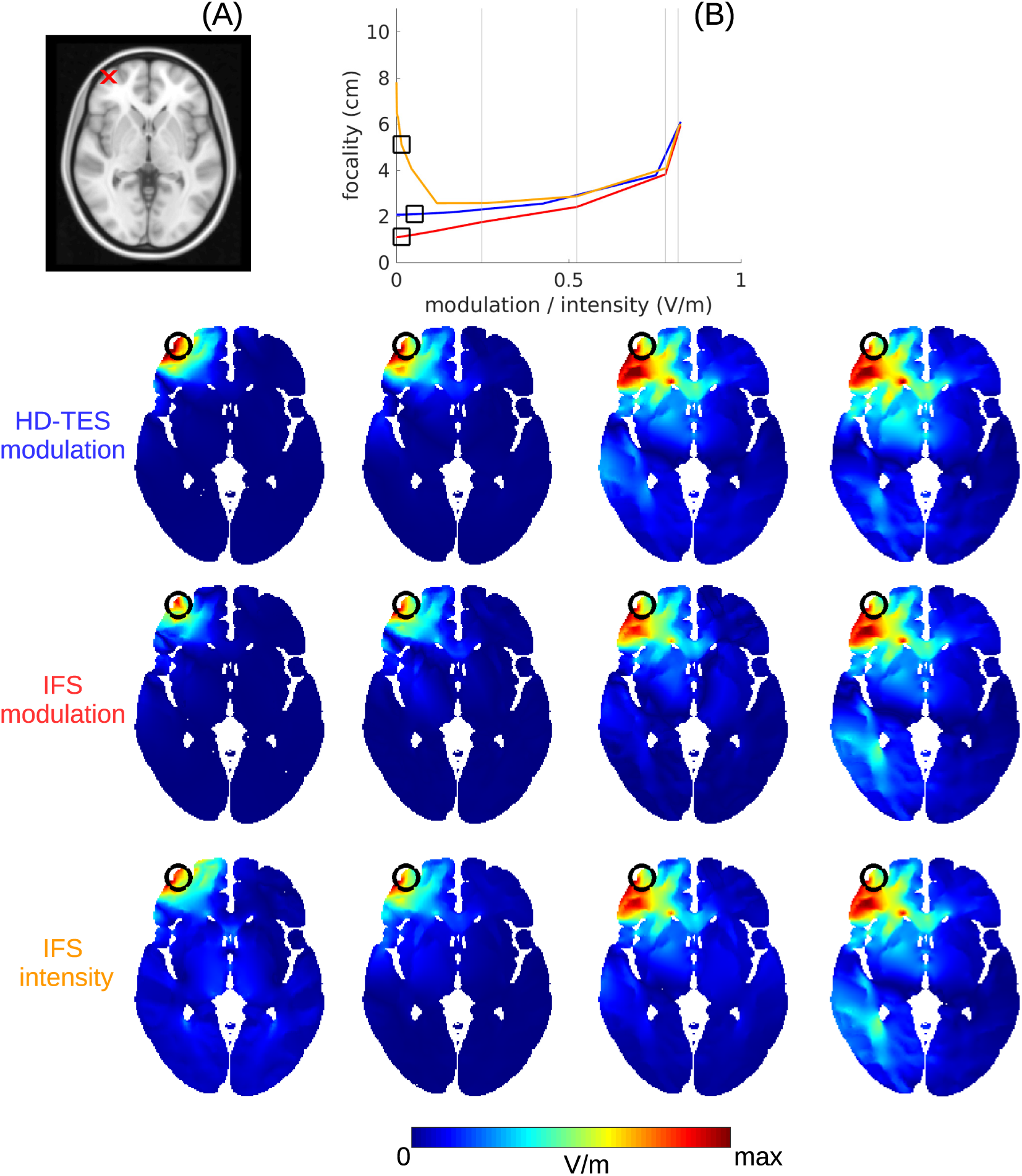
Radial component of the modulation depth / intensity under different off-target energy constraints *P*_*max*_ for the 1st target (indicated by the red x in panel (A)). The four columns of heatmaps correspond to the four gray vertical lines in panel (B), which is taken from Figure 3 C1, with *P*_*max*_ increasing from the left column to the right. The three rows of heatmaps show the HD-TES modulation depth, the IFS modulation, and the IFS intensity (text labels color-coded to correspond to the curves in panel (B).

**Figure S3:**
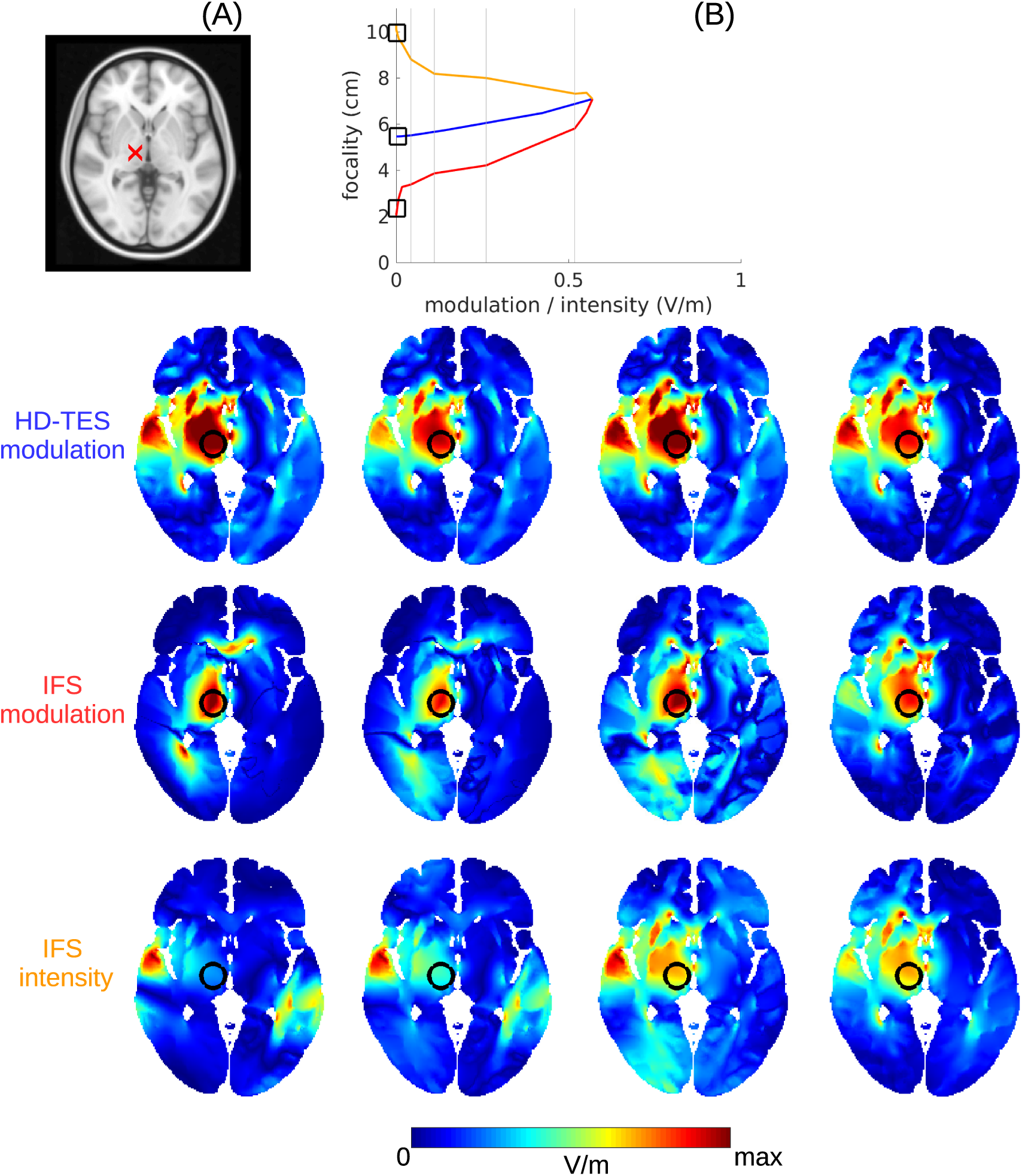
Same as in previous figure.

**Figure S4:**
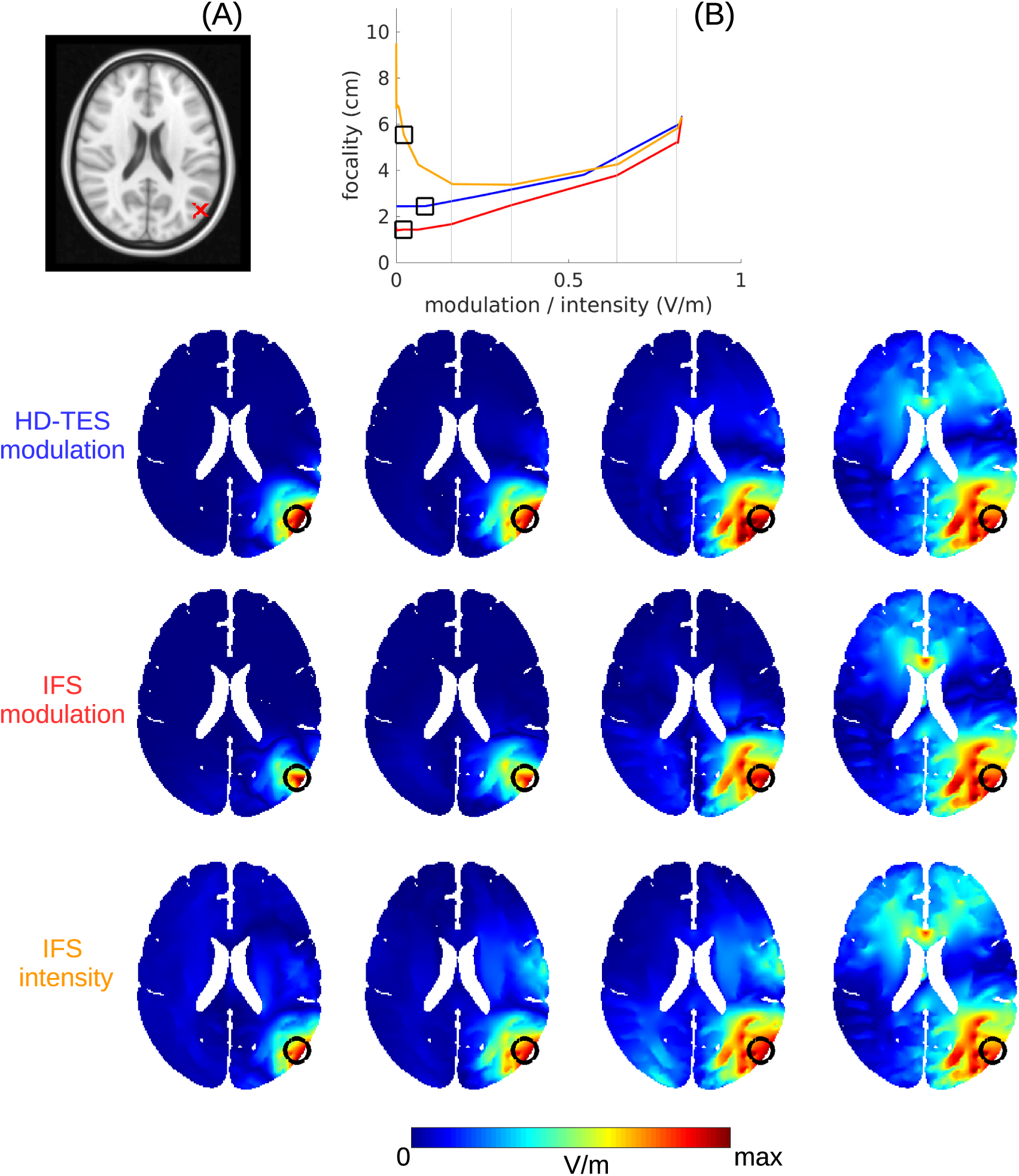
Same as in previous figure.

**Figure S5:**
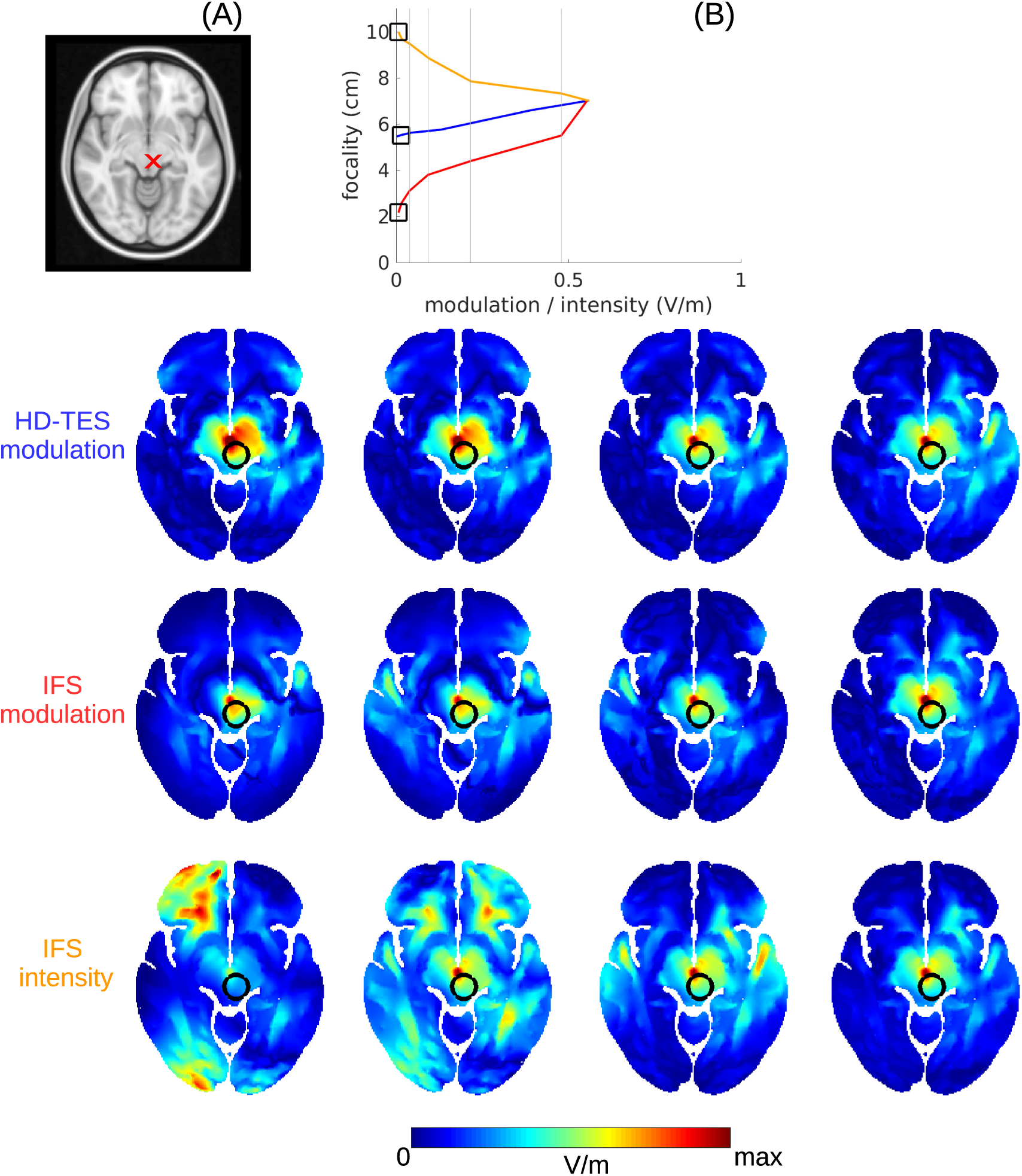
Same as in previous figure.

